## Supplementary Material for "Brain Dissection: fMRI-trained Networks Reveal Spatial Selectivity in the Processing of Natural Images"

### S1 Overview

Section S2 contains more details of the methods described in the main paper. Section S3 provides additional results from the experiments.

We include an interactive project page, which includes exploratory brain flatmaps for each of our dissection measures, and code for training and evaluating the networks. Please visit the following link to access the project page: https://brain-dissection.github.io/

### S2 Implementation details

#### S2.1 Network Architecture and Training

Our architecture and training largely follow Khosla et al. [1], except we use regular convolutions instead of E(2)-steerable convolutions. We found that using regular convolutions instead of E(2)-steerable convolutions does not significantly affect dissection performance while providing significantly improved training speed and memory efficiency. We train a separate convolutional neural network (CNN) for each ROI sub-region shared across all 8 subjects. The CNN consists of four "blocks", each with two convolutional layers followed by average pooling. We use batch normalization and early stopping. For each voxel in the ROI sub-region (across for all subjects), we employ a linear readout model on top of the feature space to predict the responses of individual voxels in a specific brain region. The linear readout is factorized into spatial and feature dimensions following [2]. This allows us to separate image spatial tuning (the "where") from feature tuning (the "what"). The spatial features have been shown to correlate with the population receptive fields (pRFs) of the voxels [3]. We train with a mean squared error loss between the predicted voxel response and the true voxel response. During the dissection procedure, we only use the feature tuning. We train all models on a single Nvidia GeForce RTX 3090 GPU. The number of parameters in the CNN is 784697 and the number of parameters in the transformer network (see next section) is 7053009.

#### S2.2 Training & Validation Curves

We provide encoding model training and validation curves for PPA, OPA and RSC functionally-defined ROIs in Figure S1 and Figure S2. We utilize early stopping to terminate training and obtain an evaluation checkpoint by checking if the validation performance (Pearson correlation) is lower than the highest validation performance for 10 epochs.

37th Conference on Neural Information Processing Systems (NeurIPS 2023).

#### S2.3 Additional Evaluation Details

For network dissection, we use the code repository from Bau et al. [4]. We use pycortex to visualize voxel data [5]. Data are projected onto 2D brain flatmaps from 3D voxel data using trilinear interpolation for continuous data (e.g., depth) and nearest interpolation for discrete data (e.g., category).

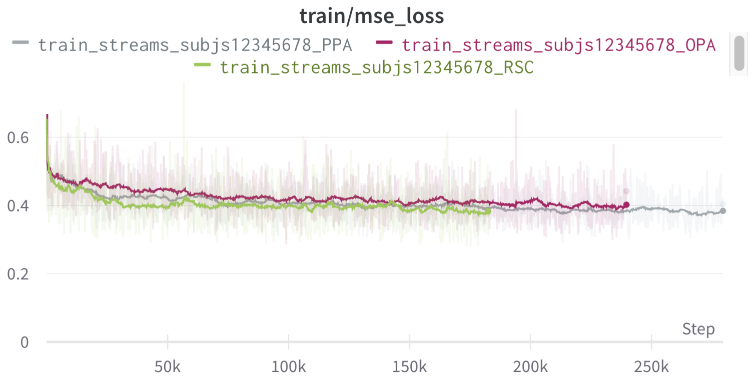


Figure S1: Mean squared error training curves for encoding models for PPA (gray), OPA (pink), and RSC (green) ROIs. X-axis is number of training steps.

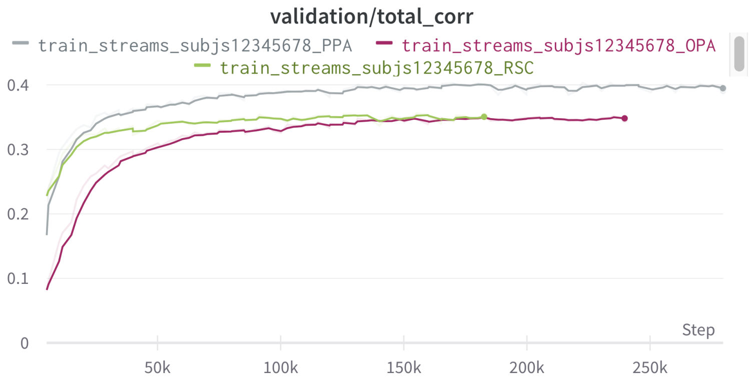


Figure S2: Pearson Correlation validation curves for encoding models for PPA (gray), OPA (pink), and RSC (green) ROIs. X-axis is number of training steps. Training was terminated using early stopping based on the validation curves.

### S3 Additional Results

#### S3.1 Performance of Response-Optimized and Baseline Models

We have provided the Pearson correlation coefficient plotted on a flatmap for all ROIs on a held-out test set of NSD data in Figure S3 for example Subject S1. Additionally, we compare our model's mean Pearson correlation to the features from Lescroart and Gallant [6] and ImageNet task-optimized features [7] on held-out NSD brain data using Pearson correlation. We the features fit to brain data via Ridge regression. We computed the features in Lescroart and Gallant [6] for NSD images using the estimation networks. As shown in Figure S3, our brain dissection model demonstrates a significant improvement, nearly 2x across scene ROIs, in its alignment with the brain responses compared to the baseline features from [6]. Our model also aligns in performance with AlexNet ImageNet features, even though the AlexNet network having a significantly larger parameter size and trained on significantly more images.

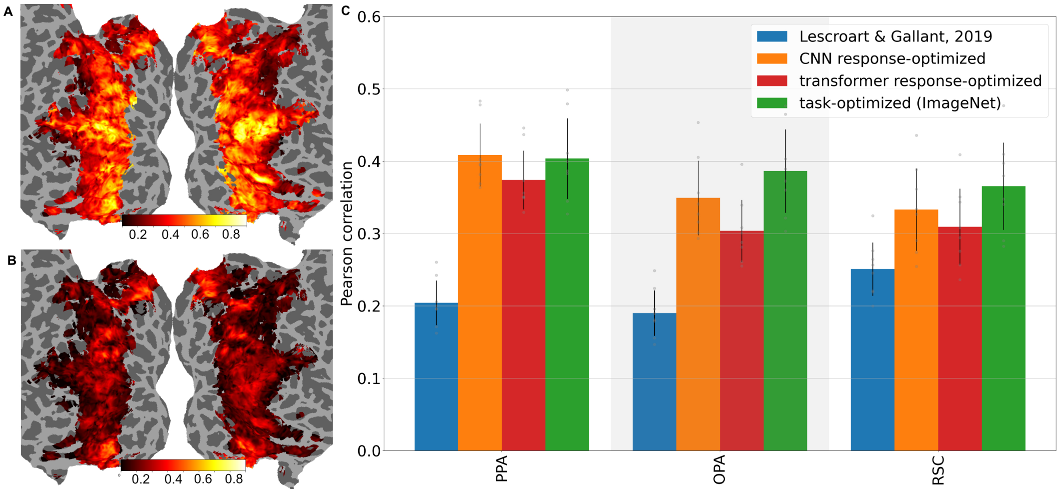


Figure S3: **A.** Pearson correlation coefficient plotted on a flatmap for all ROIs on a held-out test set of NSD data for example Subject S1 for our response-optimized networks. **B.** Same as **A** but for the features of Lescroart and Gallant [6]. **C.** Mean Pearson correlation for our model, the features from Lescroart and Gallant [6], and the ImageNet task-optimized features [7] on held-out NSD brain data using Pearson correlation for the scene-selective ROIs.

#### S3.2 Results using Different Architectures and Interpretability Methods

We integrated both gradCAM [8] and raw attention techniques [9]. The former utilizes input features combined with network gradients, while the latter leverages attention scores from transformer architectures. As shown in Figure S4, our results affirm consistency across diverse interpretability methods and network architectures.

#### S3.3 GQA Data Visualization

We show example images and annotations for the GQA dataset in Figure S5. We used Segment Anything [10] to obtain the segmentation masks from the GQA box annotations.

#### S3.4 Histograms of the Spatial Measures

We show the distribution of voxel preferences for each high-level visual ROI in Figure S6 and for the scene ROIs in Figure S7. We observe that subjects are largely consistent in their distributions for each ROI. We also observe that some ROIs and measures are multi-modal, such as in several measures in Medial and Gaussian curvature in Ventro-lateral and Lateral. This may indicate a further specialization of voxel clusters in these areas.

#### S3.5 Places365 Evaluation

In addition to the GQA dataset, we evaluate for category selectivity in the Places365 dataset, where segmentation masks are obtained using the Unified Perceptual Parsing image segmentation network [11] previously trained on 20,000 scene-centric images from the ADE20k dataset [12]. We report median IOU (mean ± standard deviation across 8 subjects) in Figure S8.

#### S3.6 Unit Visualizations

We include additional single-voxel visualizations for RSC, OPA, and PPA in Figure S9, Figure S10, and Figure S11, respectively.

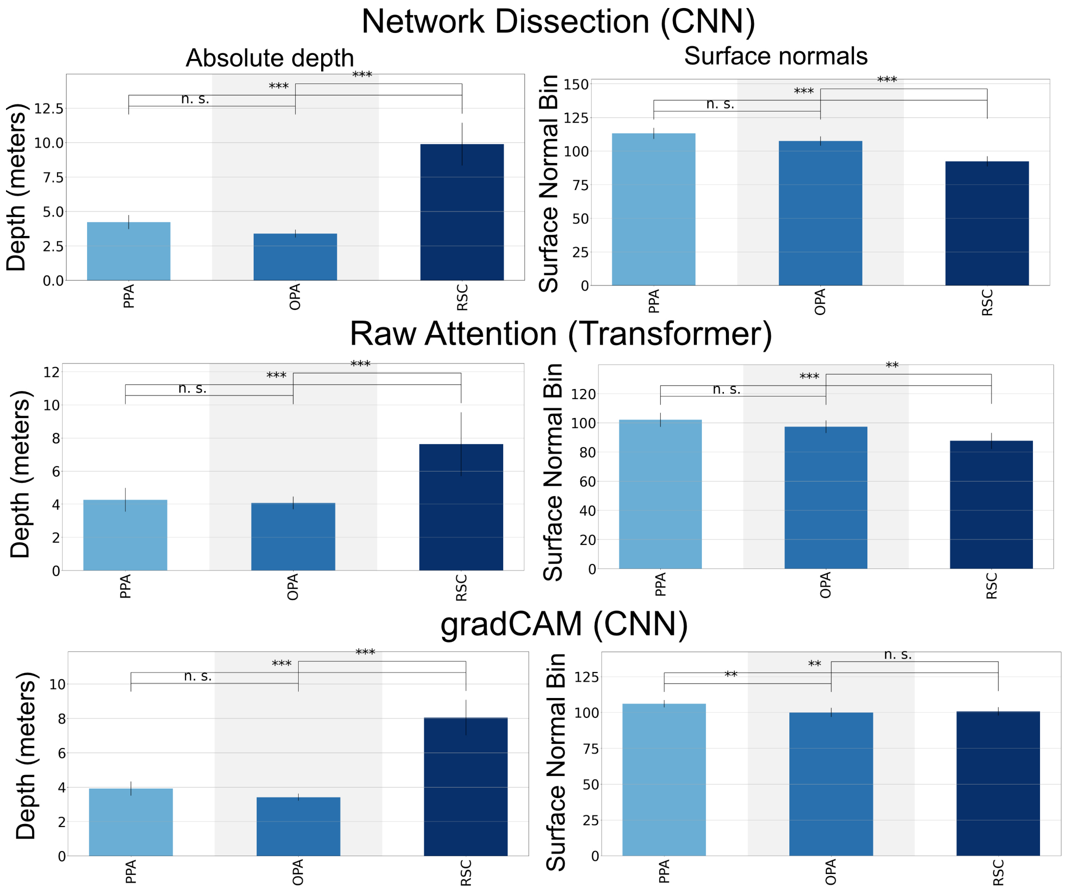


Figure S4: Depth and surface normal results for the Network Dissection [4], gradCAM [8] and raw attention techniques [9] for the scene-selective ROIs.

#### S3.7 Brain Flatmaps

In this section, we include brain flatmaps for all eight subjects for metric depth (Figure S12), relative depth (Figure S13), surface normals (Figure S14), Guassian curvature (Figure S15), and shading (Figure S16).

#### S3.8 WordCloud Visualizations

We offer a WordCloud representation to visualize the top 20 categories that an ROI selects for. In this visualization, the category size represents the magnitude of the median IOU. We show WordClouds for all high-level visual ROIs and scene ROIs for the GQA dataset (Figure S17) and the Places365 dataset (Figure S18).

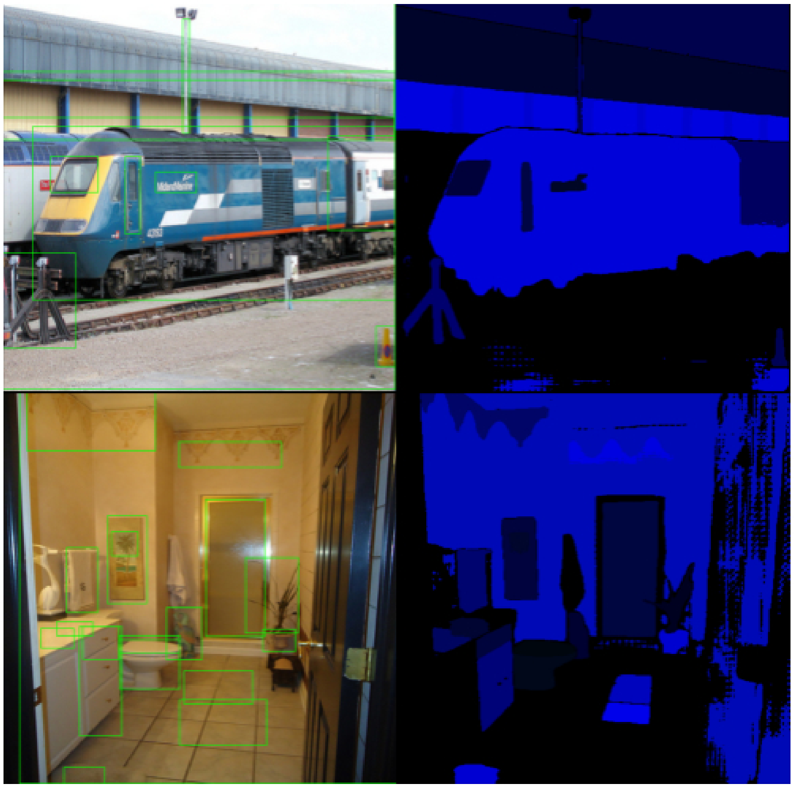

Figure S5: GQA image with bounding box annotations (left). Segment Anything [10] output given the box annotations as input (right).

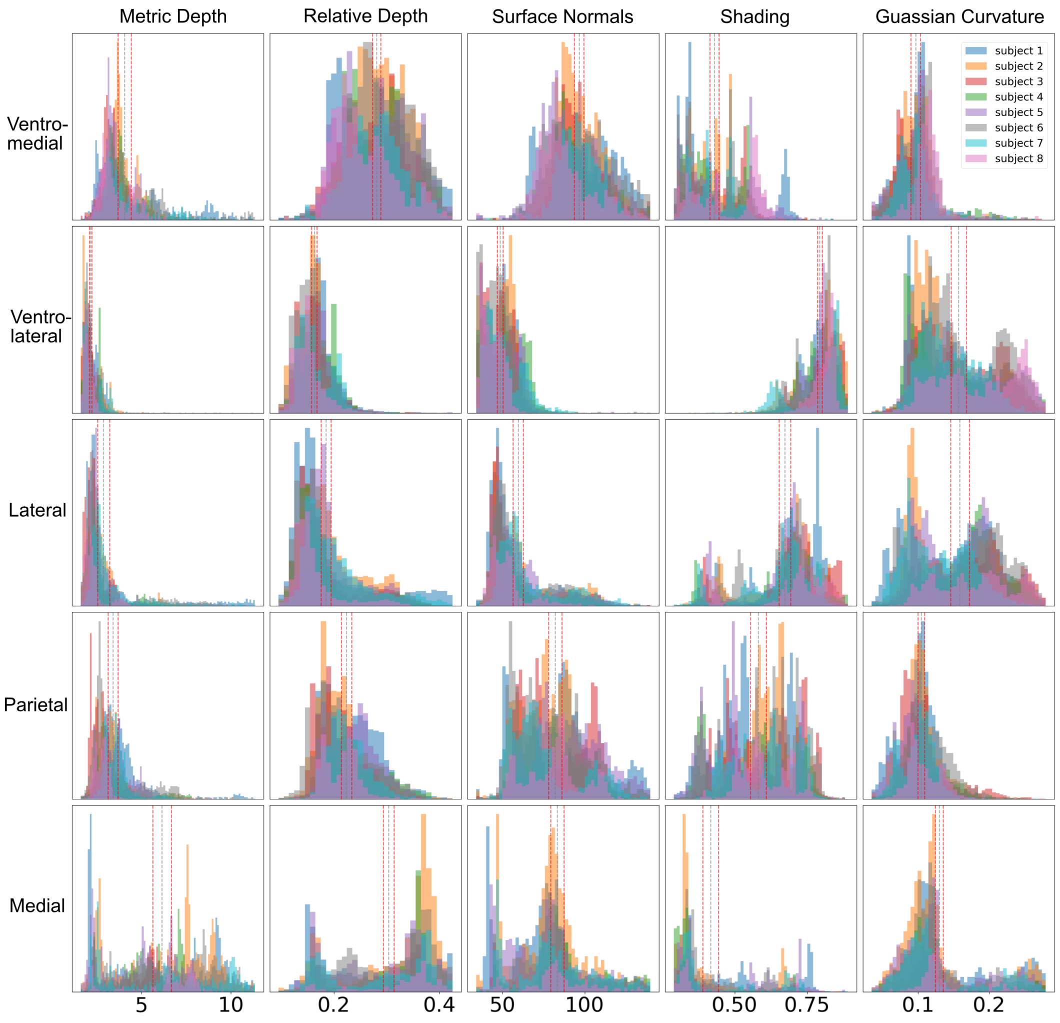


Figure S6: Histogram representation of each measure for each high-level visual ROI. Each color represents a different subject. Gray vertical dotted lines indicate the grand mean, and red vertical dotted lines indicate the 95% confidence intervals.

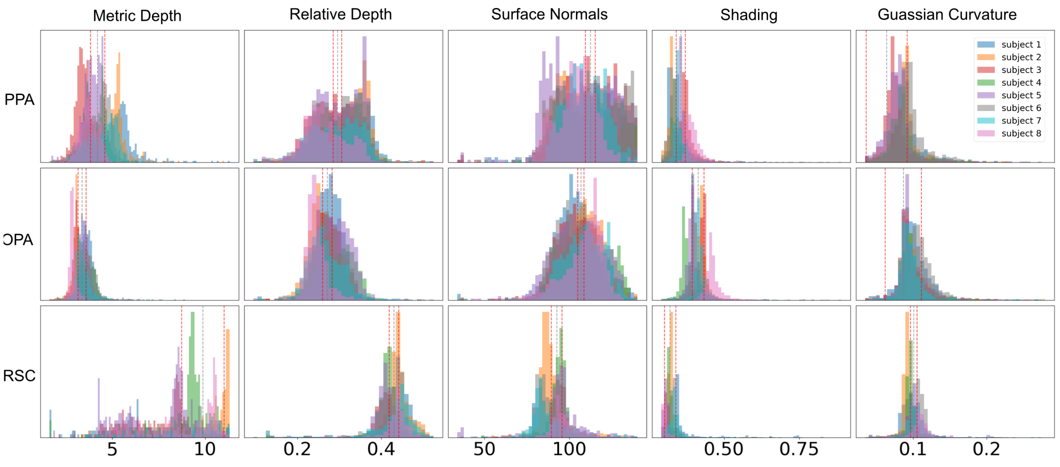


Figure S7: Histogram representation of each measure for each scene ROI. Each color represents a different subject. Gray vertical dotted lines indicate the grand mean, and red vertical dotted lines indicate the 95% confidence intervals.

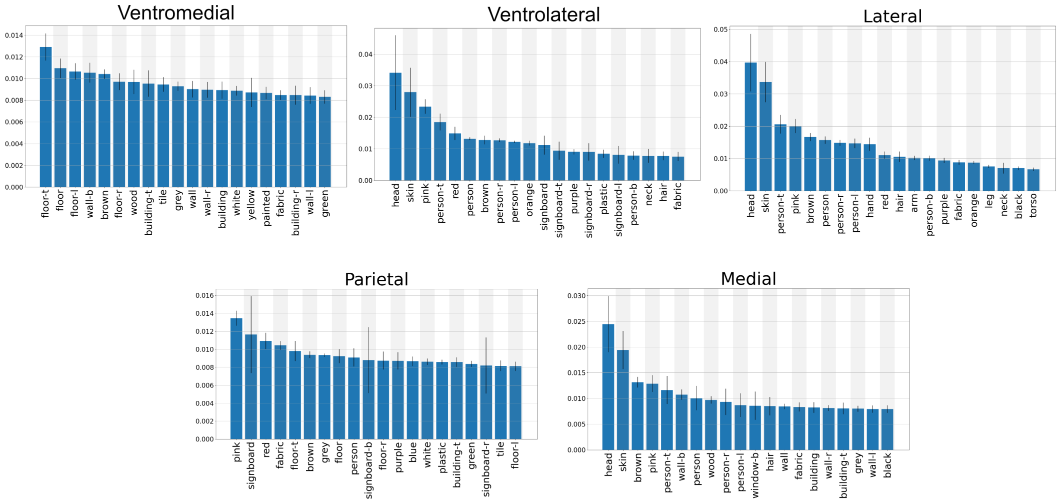


Figure S8: The median IOU (mean ± standard deviation across 8 subjects) for the top 20 concepts for each high-level visual ROI for the places365 dataset.

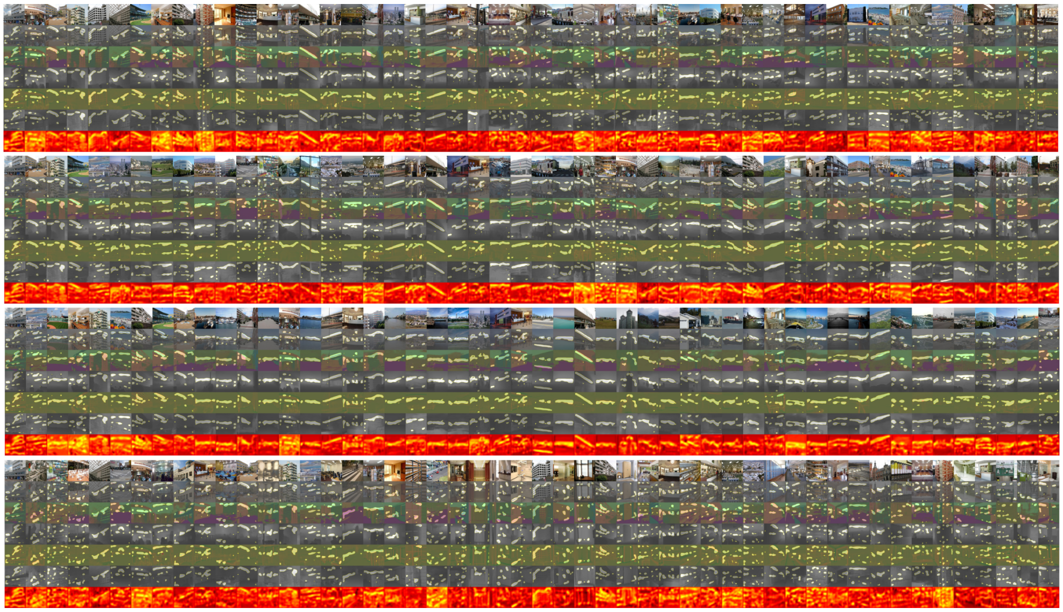

Figure S9: Unit visualization plots for four additional RSC units aligned with the mean depth selectivity of the ROI (note images are down-sampled to save memory). Each unit displays its top 50 images by predicted response for that unit. For the 50 images, we show 7 sets of images (by row - top to bottom): 1. original RGB; 2. Dissection mask overlaid on the RGB image; 3. Dissection mask overlaid on the surface normal map; 4. Dissection mask overlaid on the depth map; 5. Dissection mask overlaid on the Gaussian curvature map; 6. Dissection mask overlaid on the shading map; 7. heatmap visualization of the voxel feature map for the image.

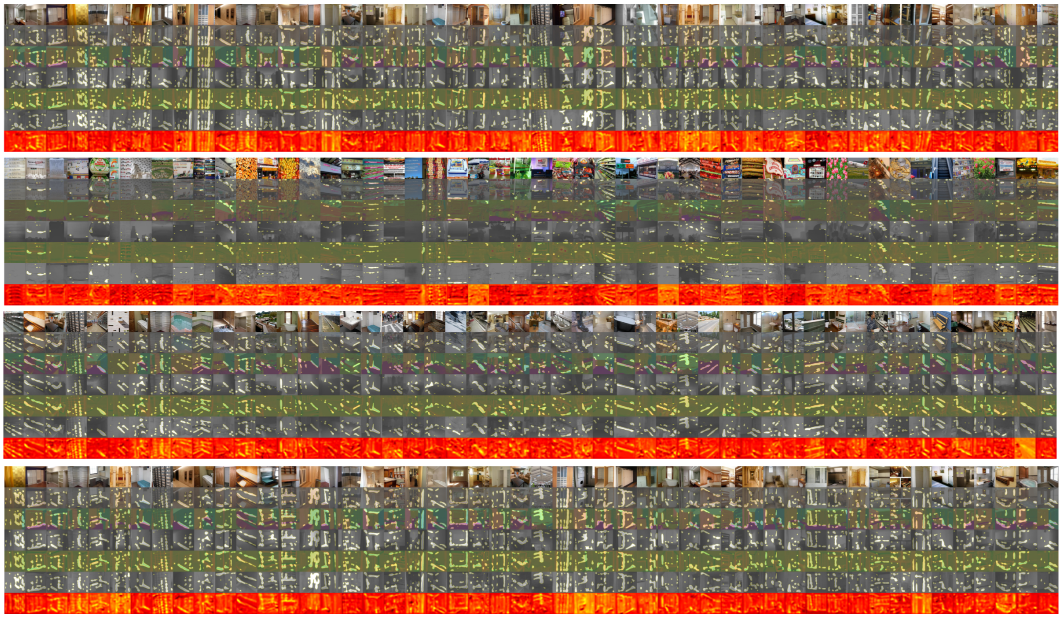

Figure S10: Unit visualization plots for four additional OPA units aligned with the mean depth selectivity of the ROI (note that images are down-sampled to save memory). Same format as Figure S9.

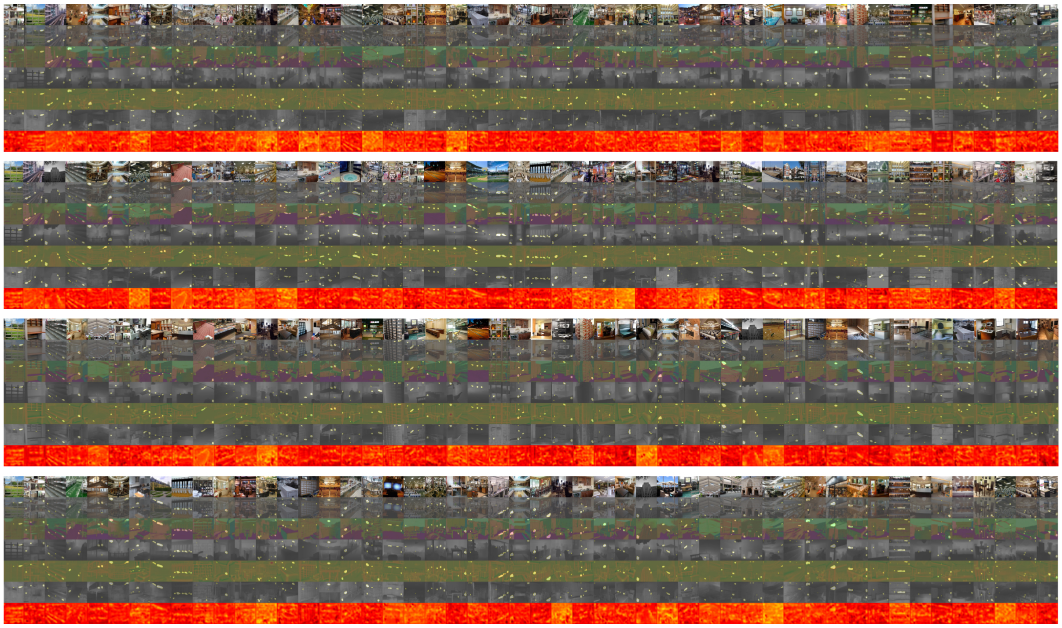

Figure S11: Unit visualization plots for four additional PPA units aligned with the mean depth selectivity of the ROI (note that images are down-sampled to save memory). Same format as Figure S9.

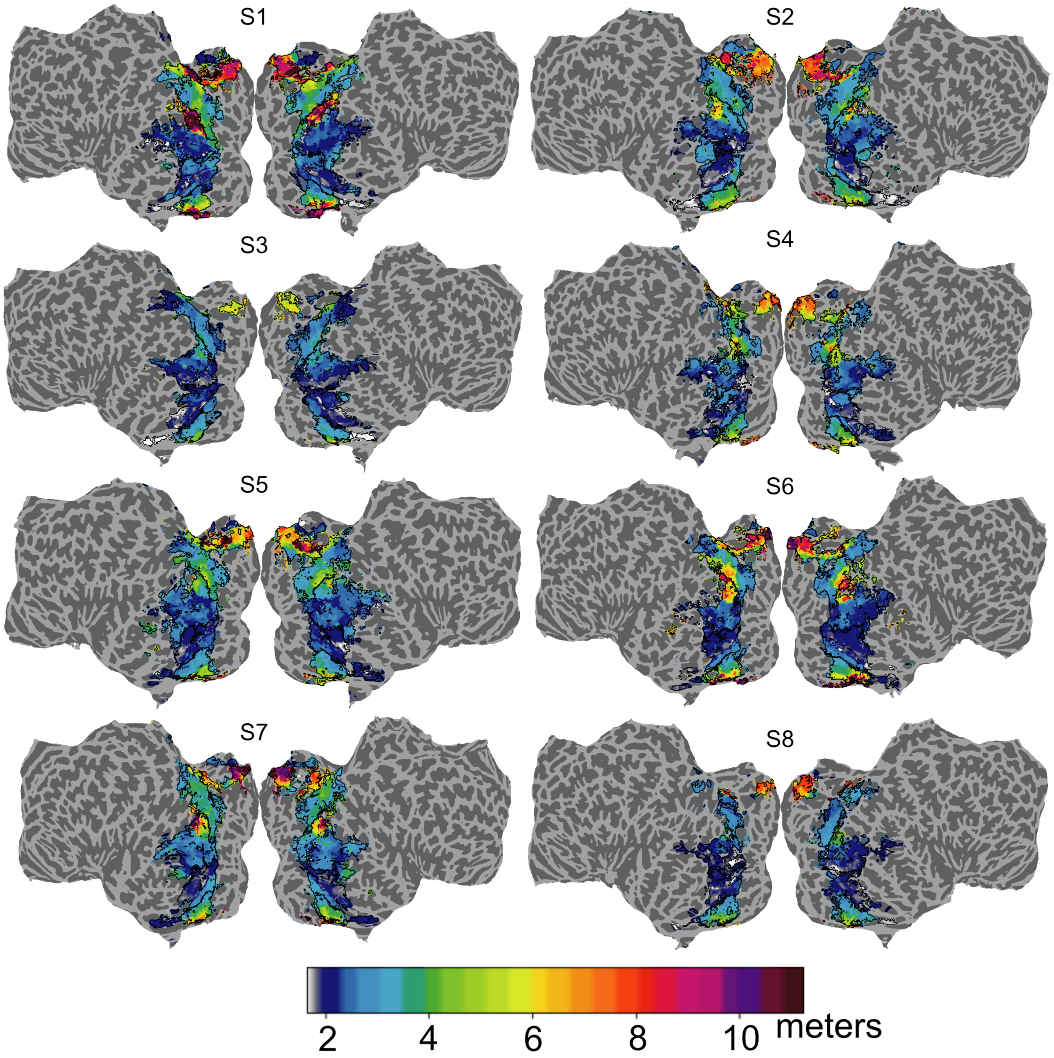


Figure S12: Metric depth on flatmaps colored by selectivity per voxel for subjects S1-S8.

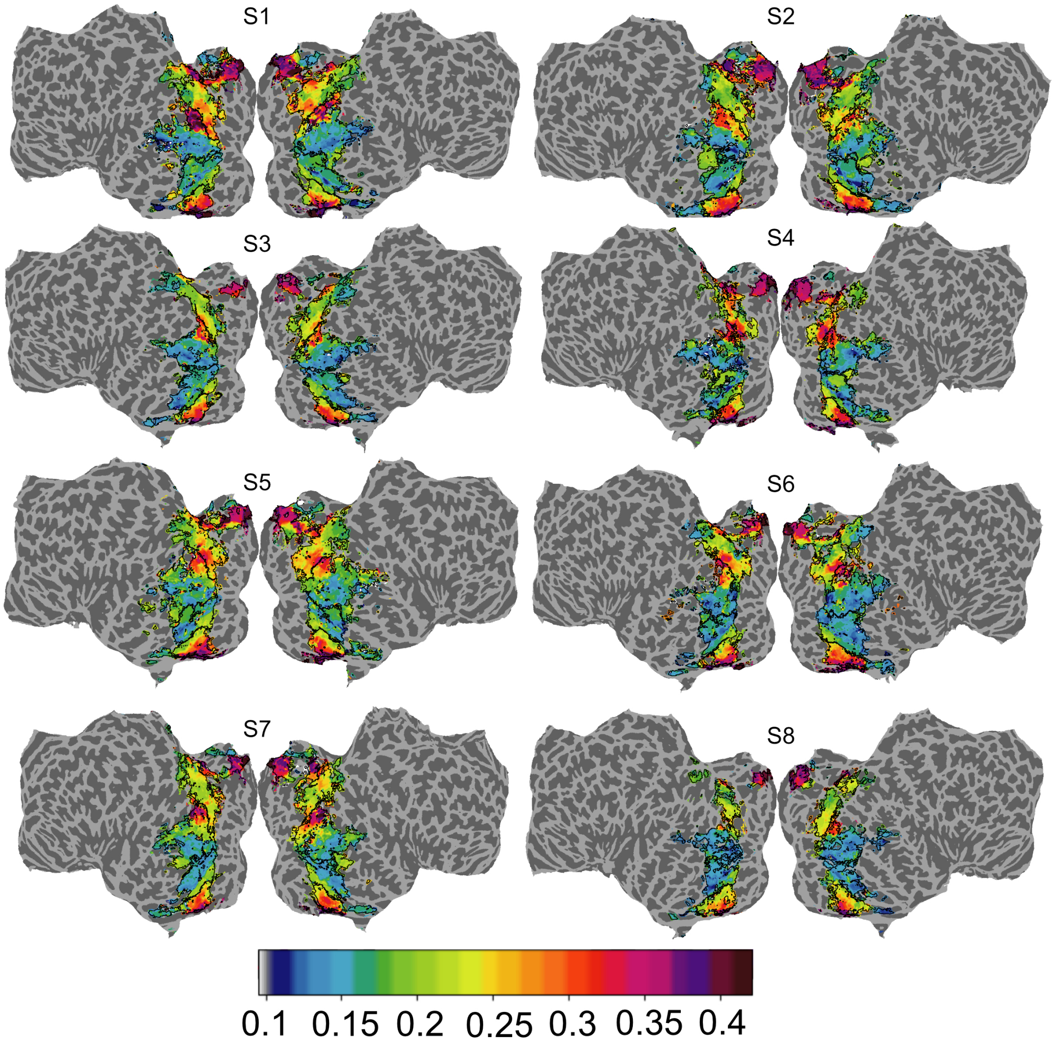


Figure S13: Relative depth on flatmaps colored by selectivity per voxel for S1-S8.

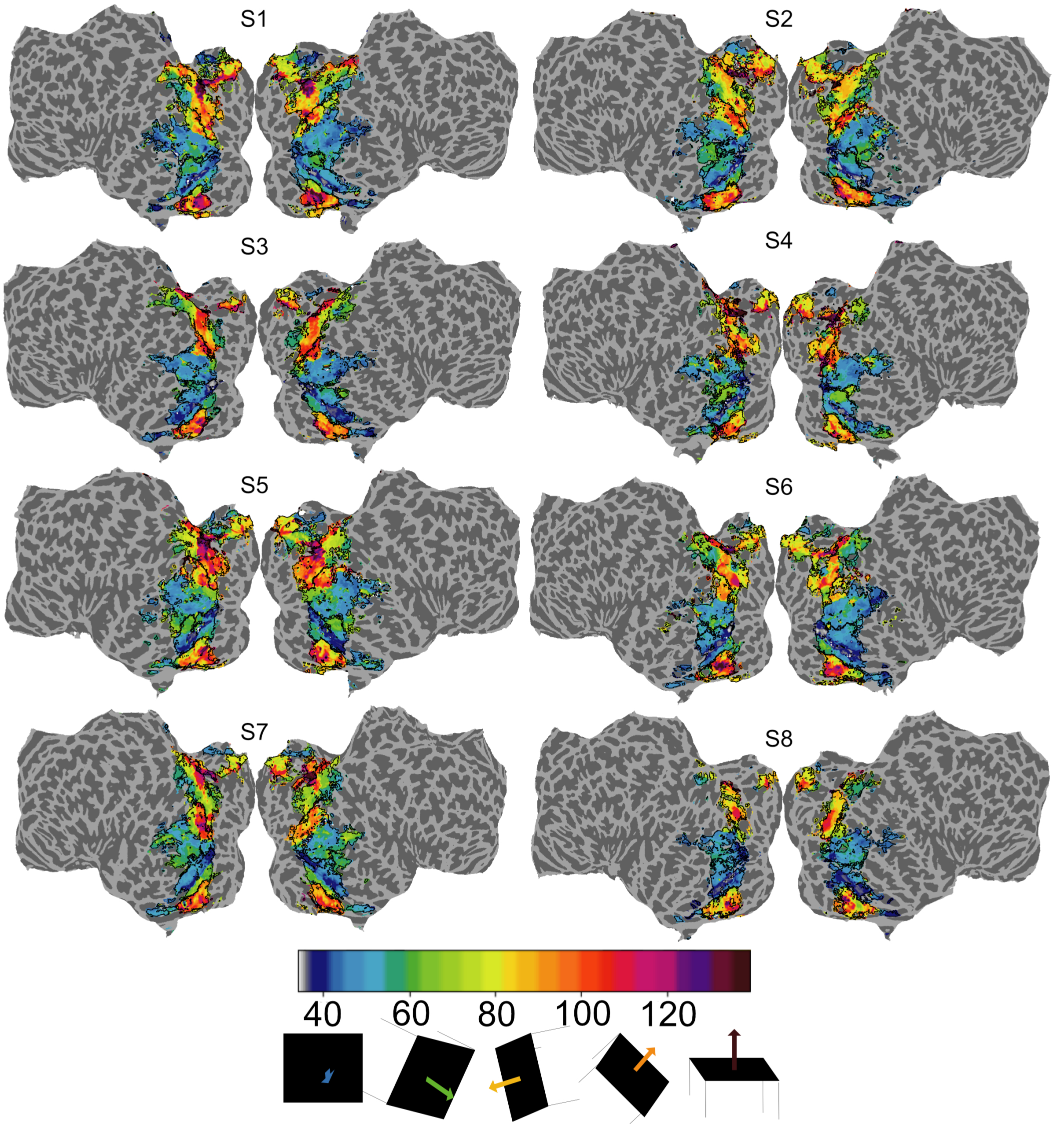


Figure S14: Surface normals on flatmaps colored by selectivity per voxel for S1-S8.

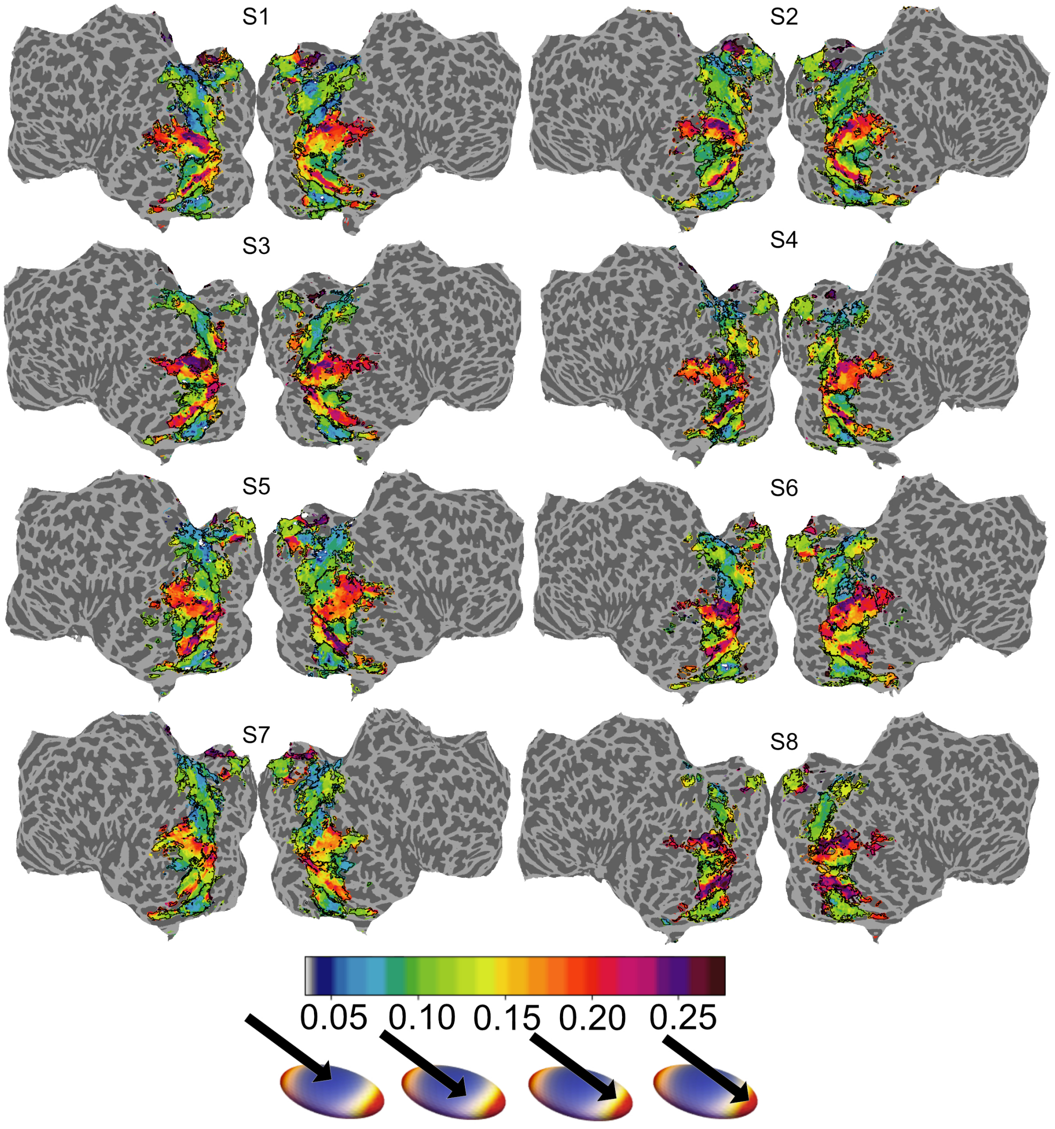


Figure S15: Guassian curvature on flatmaps colored by selectivity per voxel for S1-S8.

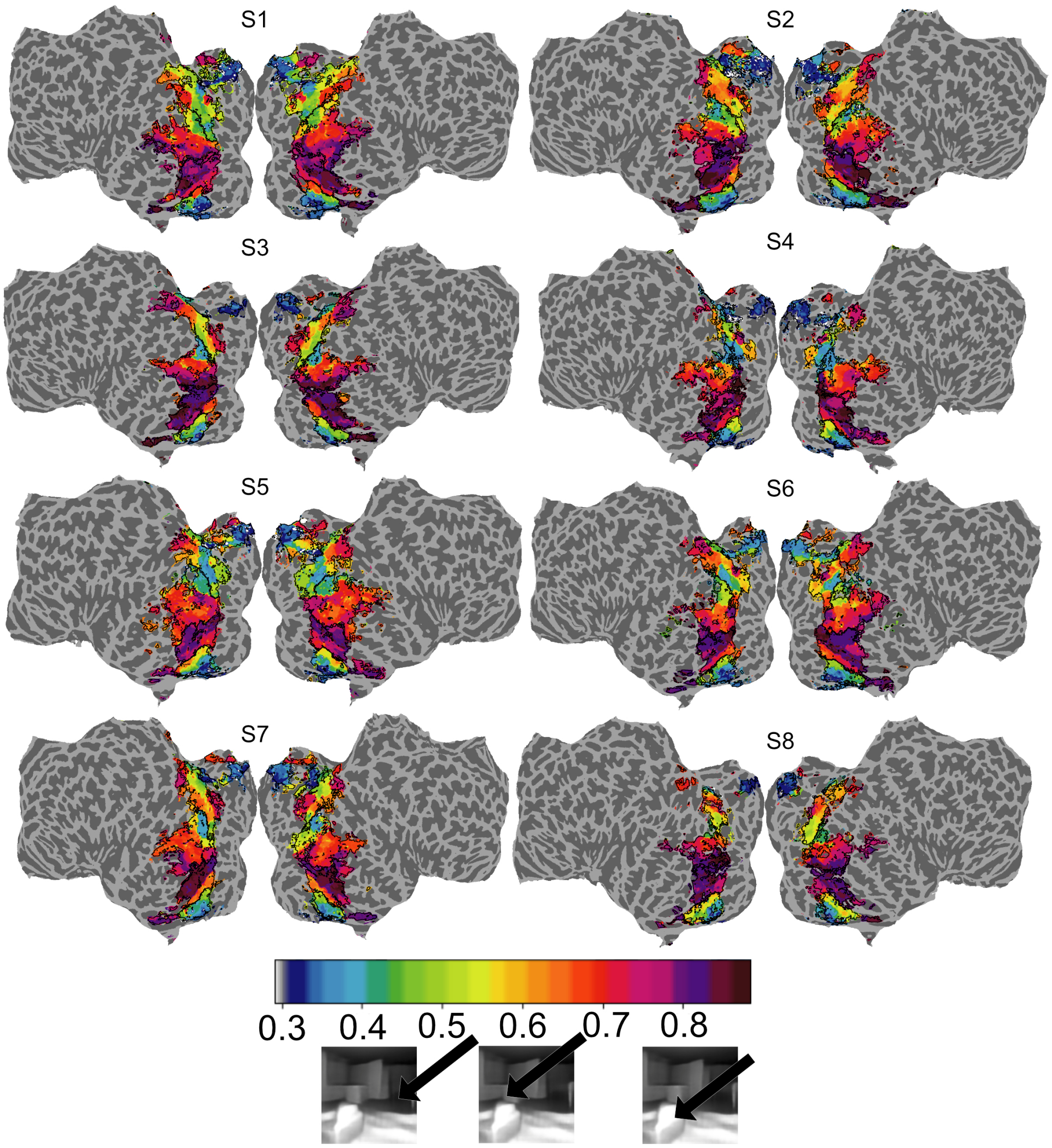


Figure S16: Shading on flatmaps colored by selectivity per voxel for S1-S8.

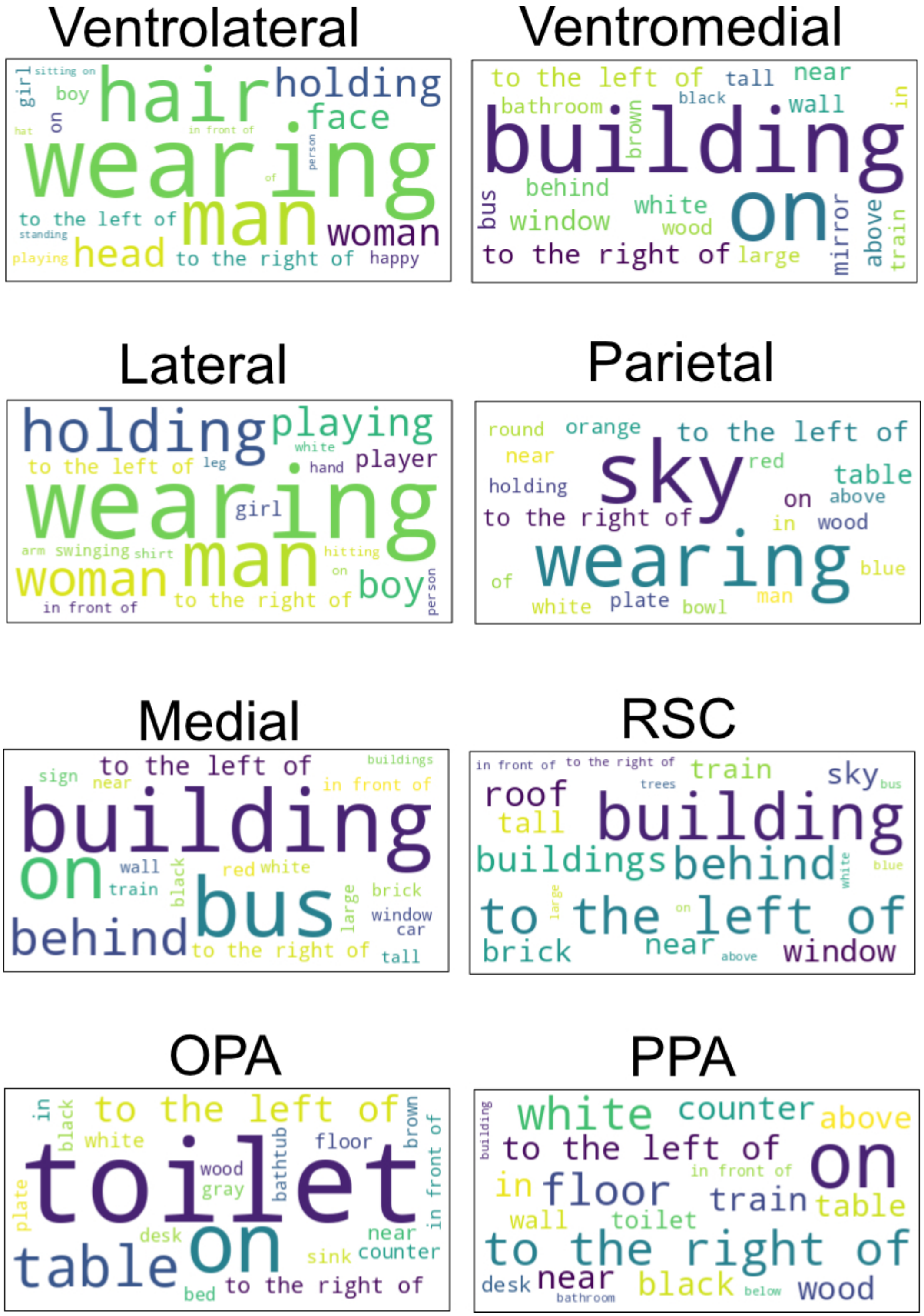


Figure S17: WordCloud for top 20 categories that an ROI selects for in the GQA dataset. In this visualization, the category size represents the magnitude of the median IOU

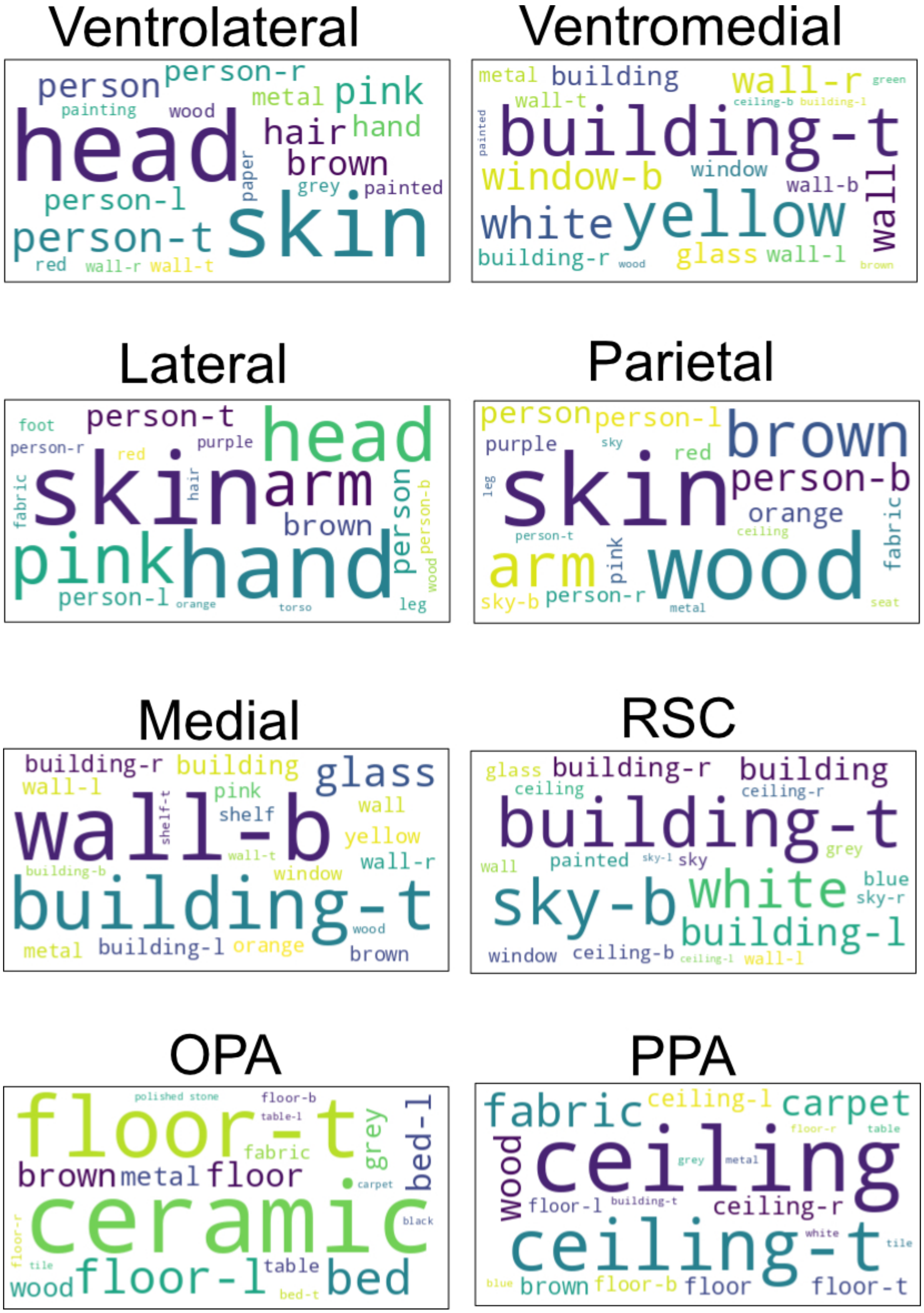


Figure S18: WordCloud for top 20 categories that an ROI selects for in the Places365 dataset. In this visualization, the category size represents the magnitude of the median IOU

[2] David Klindt, Alexander S Ecker, Thomas Euler, and Matthias Bethge. Neural system identification for large populations separating "what" and "where". *Advances in Neural Information Processing Systems*, 30, 2017.

[3] Meenakshi Khosla, Keith Jamison, Amy Kuceyeski, and Mert Sabuncu. Characterizing the ventral visual stream with response-optimized neural encoding models. *Advances in Neural Information Processing Systems*, 35:9389–9402, 2022.

[4] David Bau, Bolei Zhou, Aditya Khosla, Aude Oliva, and Antonio Torralba. Network dissection: Quantifying interpretability of deep visual representations. In *Proceedings of the IEEE conference on computer vision and pattern recognition*, pages 6541–6549, 2017.

[5] James S Gao, Alexander G Huth, Mark D Lescroart, and Jack L Gallant. Pycortex: an interactive surface visualizer for fmri. *Frontiers in neuroinformatics*, page 23, 2015.

[6] Mark D Lescroart and Jack L Gallant. Human scene-selective areas represent 3d configurations of surfaces. *Neuron*, 101(1):178–192, 2019.

[7] Alex Krizhevsky, Ilya Sutskever, and Geoffrey E. Hinton. Imagenet classification with deep convolutional neural networks. In *Proceedings of the 25th International Conference on Neural Information Processing Systems - Volume 1*, NIPS'12, page 1097–1105, Red Hook, NY, USA, 2012. Curran Associates Inc.

[8] Ramprasaath R Selvaraju, Michael Cogswell, Abhishek Das, Ramakrishna Vedantam, Devi Parikh, and Dhruv Batra. Grad-cam: Visual explanations from deep networks via gradient-based localization. In *Proceedings of the IEEE international conference on computer vision*, pages 618–626, 2017.

[9] Mathilde Caron, Hugo Touvron, Ishan Misra, Hervé Jégou, Julien Mairal, Piotr Bojanowski, and Armand Joulin. Emerging properties in self-supervised vision transformers. In *Proceedings of the International Conference on Computer Vision (ICCV)*, 2021.

[10] Alexander Kirillov, Eric Mintun, Nikhila Ravi, Hanzi Mao, Chloe Rolland, Laura Gustafson, Tete Xiao, Spencer Whitehead, Alexander C Berg, Wan-Yen Lo, et al. Segment anything. *arXiv preprint arXiv:2304.02643*, 2023.

[11] Tete Xiao, Yingcheng Liu, Bolei Zhou, Yuning Jiang, and Jian Sun. Unified perceptual parsing for scene understanding. In *Proceedings of the European conference on computer vision (ECCV)*, pages 418–434, 2018.

[12] Bolei Zhou, Hang Zhao, Xavier Puig, Sanja Fidler, Adela Barriuso, and Antonio Torralba. Scene parsing through ade20k dataset. In *Proceedings of the IEEE conference on computer vision and pattern recognition*, pages 633–641, 2017.
